# Rapid Identification of Metal Resistance Genes Using an Enhanced ResNet Deep Learning Model Trained on a largely Expanded BacMet-Based Database

**DOI:** 10.1101/2025.07.04.663169

**Authors:** Jiale Chen, Xianli Gao, Chunhua Zhang, Ying Ge

## Abstract

Heavy metal pollution poses significant risks to both the environment and public health. Effective management requires not only reducing contaminants but also understanding microbial adaptation, which could be achieved through the comprehensive identification and classification of metal resistance genes. This study expanded the existing BacMet database by incorporating 1,219,137 unique amino acid sequences through BLASTp analysis, thereby increasing the number of metal resistance-related acid sequences by more than **1,600-fold** compared to the 753 sequences included in the previous version. We employed various deep learning models for our proposed multi-label framework which could effectively overcome the well-recognized challenging issue of strict classification among metal translocating proteins such as CopA, ZntA, SilA, CadA, and CzcA due to their high sequence similarity and overlapping metal specificities, including AlexNet, VGG, GoogleNet, ResNet, BERT, ViT, and Mamba2, with ResNet demonstrating superior performance in terms of accuracy, robustness, and computational efficiency. The ResNet model achieved a Jaccard score of 98.91%, significantly higher than BLASTp (98.19%) and DIAMOND (98.16%), and was approximately 6,500 times faster in inference speed than these traditional alignment-based methods. Furthermore, the predictive performance of the model and the reliability of the expanded gene library were experimentally validated through overexpression and metal resistance assays of selected genes. Additionally, we developed the Predicting Metal Resistance Amino Acid Sequences (PMRAAS, https://s3.v100.vip:16165) website, facilitating online predictions of gene metal resistance. Our findings provided deeper insights into microbial adaptation mechanisms in metal-polluted environments and offered a robust tool for advancing heavy metal pollution management through enhanced bioremediation strategies.

## 1 Introduction

Heavy metal contamination poses a serious threat to the environment and public health, affecting quality of soil, water, and biodiversity (Briffa et al., 2020; Das et al., 2023). Long-term exposure can result in acute and chronic health issues, particularly for children and vulnerable populations, necessitating global governance and mitigation measures(Adnan et al., 2024; Jaishankar et al., 2014). Various countries and international organizations are promoting measures to address heavy metal pollution, including the application of cleaner production technologies, pollution source control, and the development of remediation techniques(Sánchez-Castro et al., 2023). Additionally, bioremediation technologies using microorganisms and plants, have become a key research focus for mitigating heavy metal pollution(Raklami et al., 2022). Microorganisms respond to heavy metal pollution through metal resistance genes, utilizing mechanisms such as adsorption, absorption, redox reactions, and the secretion of extracellular polymeric substances (EPS) to reduce metal toxicity, enabling survival and adaptation in polluted environments, and maintaining ecological balance(Igiri et al., 2018). Understanding microbial metal resistance genes helps reveal their potential in metal pollution management, providing a theoretical foundation and practical guidance for the development of novel bioremediation technologies.

With the rapid development of high-throughput sequencing technologies, an increasing number of microbial genomes have been sequenced and made publicly available(McAdam et al., 2014). These advancements enable researchers to rapidly and comprehensively analyze the genomes of various microorganisms(Satam et al., 2023), thereby obtaining more information on metal resistance genes. While existing metal resistance gene databases such as BacMet(Pal et al., 2014) provided valuable information on metal resistance genes, they have slow data updates and limited scope. For instance, the latest update of the BacMet database was in 2018, and the amino acid sequences included in the experimental data are mostly derived from published literature, totaling only 753 sequences. These limitations highlight the need to expand the data on metal resistance genes.

As a classical gene annotation tool, BLASTp(McGinnis and Madden, 2004) has been widely used in genomic studies. However, it relies on known databases, cannot handle unknown gene sequences, and has limitations in predicting complex gene functions, with high computational costs and low efficiency. Deep learning offers significant advantages in gene annotation by autonomously extracting gene sequence features without relying on databases, capturing complex nonlinear relationships, improving annotation accuracy, and handling large-scale data to efficiently annotate new genes and complex sequences(Brandes et al., 2022; Kulmanov et al., 2018; Seo et al., 2018). Bileschi proposed the ProtCNN model, which improved the annotation accuracy of the protein family database Pfam(Bileschi et al., 2022). Similarly, deep learning-based AlphaFold2 predicted protein structures with both speed and accuracy(Jumper et al., 2021). These were based on mainstream deep learning models: AlexNet(Krizhevsky et al., 2017), VGG(Simonyan, 2014), GoogleNet(Szegedy et al., 2015), and ResNet(He et al., 2016) were suitable for image processing, with strong feature extraction capabilities; BERT(Devlin et al., 2019) and ViT(Dosovitskiy, 2020) were designed for sequence data processing, excelling at capturing contextual information; Mamba2(Dao and Gu, 2024) focused on multitask learning. Although recent architectures such as EfficientNet (Tan and Le, 2019), DenseNet (Huang et al., 2018), and Vision Transformers (ViTs) have shown promising performance in general computer vision tasks, ResNet was selected in this study based on its balance between performance, interpretability, and computational efficiency. Compared with deeper or more complex models, ResNet offers strong feature extraction capabilities through its residual connections while maintaining training stability, especially on medium-sized datasets like the BacMet-derived protein sequences. In our benchmark comparisons, ResNet outperformed or matched other models in terms of Jaccard score while offering faster inference time and requiring fewer computational resources. These attributes make ResNet particularly suitable for rapid annotation tools, where model speed, robustness, and practical deployment are critical.

To address the increasingly severe problem of heavy metal pollution, there was an urgent need for more metal resistance gene data to uncover the resistance mechanisms of different microorganisms and identify new resistance genes. This study constructed a metal resistance amino acid sequence library through BLASTp, applied deep learning models for multi-label framework (i.e., predicting the presence of multiple metal resistance traits simultaneously for a single sequence), and compared the performance of different models. The multi-label framework proposed in this paper could effectively overcome the well-recognized issue of strict classification among metal translocating proteins such as CopA, ZntA, SilA, CadA, and CzcA in traditional classification methods, due to their high sequence similarity and overlapping metal specificities. Innovative preprocessing methods, such as the removal of low-frequency amino acids, oversampling, and random amino acid replacement, are employed to improve the data quality and generalization ability of the models. These methods effectively optimized gene annotation accuracy and enhanced the model’s adaptability to complex data. This study also built upon previous research and compared the impact of different protein embedding methods on model performance. Finally, fast and accurate models were selected, which not only provide deeper insights into how microorganisms adapt to metal-polluted environments but also lay the foundation for the development of new bioremediation technologies. In general, our proposed method is more accurate, comprehensive, biologically meaningful, and theoretically grounded. It provides a deeper and more nuanced understanding of the complexity of metal resistance. As a result, based on our trained optimal model, we developed the Predicting Metal Resistance Amino Acid Sequences (PMRAAS, https://s3.v100.vip:16165) website, where users could predict the metal resistance of genes online. With the accumulation of more microbial genomic data and the expansion of databases, we would be able to comprehensively understand the distribution, function, and evolutionary patterns of metal resistance genes, thereby advancing the field of heavy metal pollution management.

## 2. Materials and methods

### 2.1 Data Construction

Based on the BacMet metal resistance gene database, we identified that the resistance genes belonged to 64 bacterial species. We then downloaded all reference amino acid sequences for these 64 bacterial species from the NCBI database, resulting in a total of 224,081 sequences. Next, we utilized the BLASTp tool to match these amino acid sequences to the metal resistance genes in the BacMet2_EXP database. The specific parameters used during the BLASTp matching process(Douglas and Shapiro, 2024) included: outfmt 6 for tabular output format, --evalue 1e-5 to filter out low-confidence hits (as lower e-value thresholds help eliminate random or spurious alignments, thereby improving the reliability of gene function assignment), -p 40 to enable parallel processing with 40 threads, --max-target-seqs 1 to retain only the top hit per query, --min-score 60 to exclude weak alignments, --id 40 to set a minimum sequence identity threshold of 40%, and --query-cover 20 to require a minimum of 20% coverage of the query sequence. These settings were chosen to balance sensitivity and specificity, ensuring robust identification of metal resistance gene candidates while minimizing false positives. The selected thresholds (evalue 1e-5, min-score 60, identity 40%, query coverage 20%) were adapted from previous studies(Douglas and Shapiro, 2024) and empirically validated in our setting to balance sequence similarity, functional relevance, and downstream model performance. This process yielded 18,088,297 matched sequences. To enhance the model’s generalization ability, we eliminated 100% identical sequences, ultimately retaining 1,219,137 unique sequences (Fig. 1).

**Figure 1.**
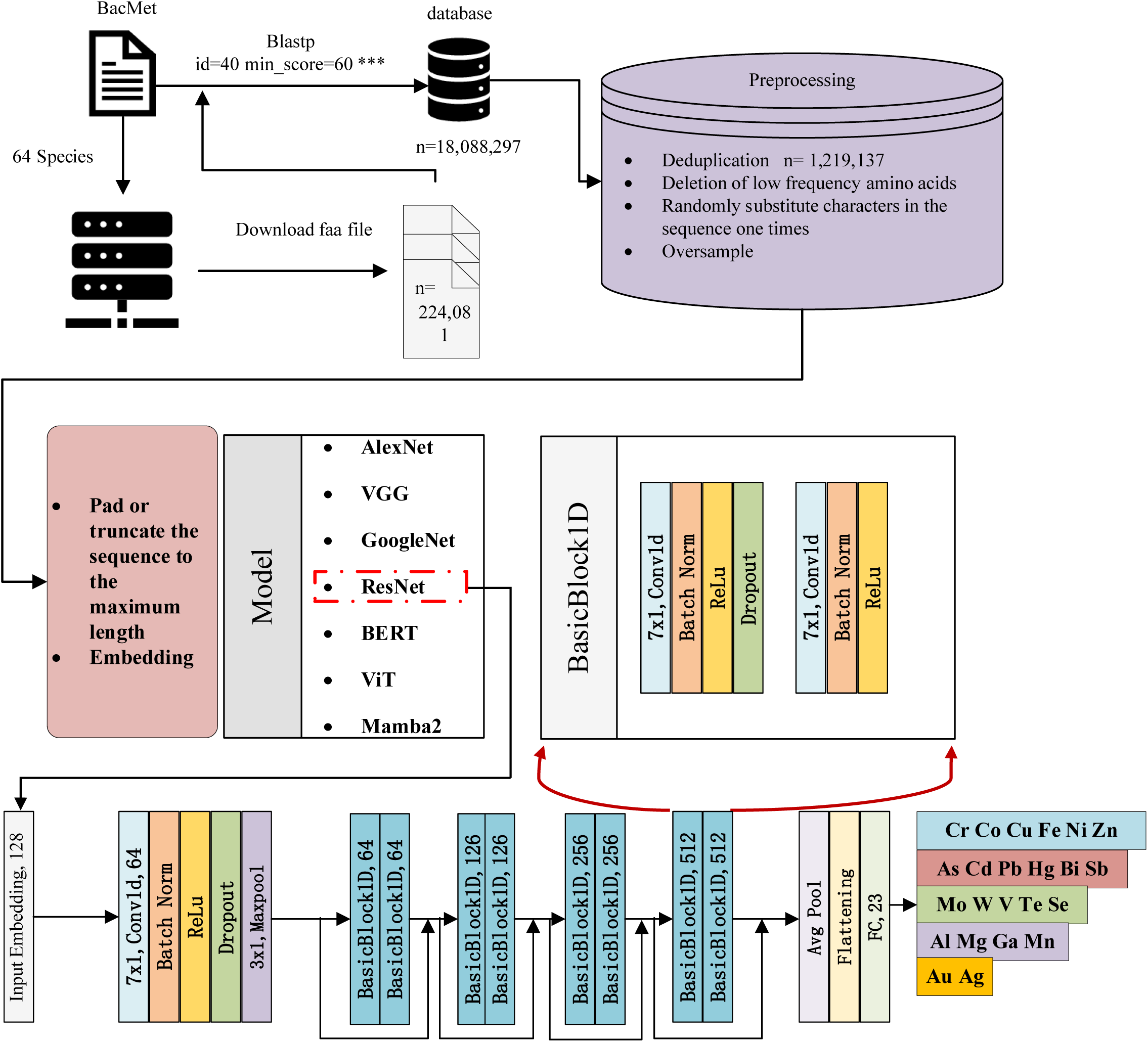
Schematic diagram of the system workflow. (1) The protein sequence files (faa) of 64 species from BacMet were downloaded from NCBI, totaling 224,081 sequences. (2) The metal resistance amino acid sequences of these 64 species, derived from BacMet, were obtained via Blastp, resulting in a total of 18,088,297 metal resistance amino acid sequences. (3) Duplicate sequences were removed, leaving 1,219,137 unique sequences. (4) Low-frequency amino acids were eliminated, and the sequences were subjected to random substitution and oversampling.(5) Based on step (4), the amino acid sequences were truncated, padded, and embedded, followed by processing through AlexNet, VGG, GoogleNet, ResNet, BERT, ViT, and Mamba2, with ResNet being selected as the optimal model. The non-redundant dataset and related code are available at https://doi.org/10.5281/zenodo.14558323.

### 2.2 Data Preprocessing and Embedding Methods

#### Data Preprocessing

Initially, we removed amino acids that occurred infrequently across all metal resistance amino acid sequences. These low-frequency amino acids were considered potential noise, which could interfere with model training and adversely affect its performance.

#### Data Augmentation

To increase the sample size, we performed random deletions, swaps, insertions, substitutions, and combinations of these operations on the amino acid sequences, analogous to image augmentation techniques used in image classification tasks. This approach facilitated the generation of additional samples and the identification of optimal combinations, thereby enhancing the model’s robustness and generalization capability.

#### Handling Sample Imbalance

To address class imbalance, both oversampling of minority classes and undersampling of majority classes were performed. Specifically, oversampling was conducted by duplicating instances from underrepresented classe(minority classes), whereas undersampling involved randomly reducing samples from overrepresented classes (majority classes). The target was to ensure that all classes had approximately equal sample sizes, thereby preventing bias during model training and improving overall performance.

#### Sequence Length Standardization

All sequences were standardized to a uniform input length. We selected the maximum length from the metal resistance amino acid sequences and set 1024 as the standard input length. Sequences shorter than 1024 were padded (typically with 0s), while those longer than 1024 were truncated, ensuring consistency in input length across all sequences.

#### Selection of Embedding Methods

To further enhance the model’s performance, we employed a variety of embedding methods, including PyTorch’s Embedding (nn.Embedding), One-Hot encoding, physicochemical embeddings(Licari et al., 2023), ProtVec(Asgari and Mofrad, 2015), ESM-1b(Rives et al., 2021), and TAPE(Rao et al., 2019). These diverse embedding techniques facilitated the capture of information at multiple levels and enriched the model’s ability to understand the characteristics of amino acid sequences.

### 2.3 Evaluation Metrics

Since the metal resistance amino acid sequences in this study could involve resistance to multiple metals, the task was classified as a multi-label classification problem. The following evaluation metrics were selected for model assessment: Jaccard score(Jaccard, 1901), F1 Macro average, Precision, Recall, and Hamming Loss.

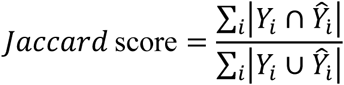

Here, *Y*_*i*_ represents the true label set, and *Ŷ_i_* denotes the predicted label set; *Σ_i_* indicates summation over all samples.

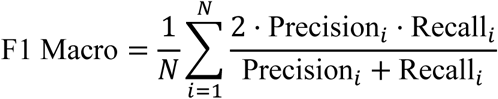

In this equation, *N* is the total number of classes; Precision_*i*_ refers to the precision of the *i*-th class, and Recall_*i*_ refers to the recall for the *i*-th class.

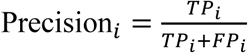

Where TP_*i*_ is the true positive count for the *i*-th class, and FP_*i*_ is the false positive count for the *i*-th class.

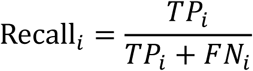

Where TP_*i*_ is the true positive count for the *i*-th class, and FP_*i*_ is the false positive count for the *i*-th class.

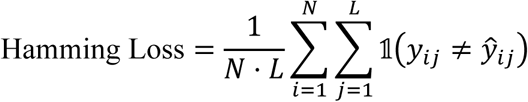

In this case, *N* represents the number of samples, *L* is the number of labels, *y*_*i*_ refers to the true value (0 or 1) of the *j*=-th label for the *i*-th sample, and *ŷ*_*ij*_ refers to the predicted value (0 or 1) of the *j*-th label for the *i*-th sample. The indicator function 𝟙 takes the value 1 if *y*_*ij*_ ≠ *ŷ*_*ij*_, otherwise, it is 0.

In this study, the task involved processing one-dimensional data, rather than the traditional two-dimensional image classification tasks. Therefore, all models were adjusted to accommodate one-dimensional data input. The core objective of these models was multi-label classification, achieved by effectively extracting temporal features for classification. To efficiently capture relevant patterns in amino acid sequences, we employed 1D convolutional models such as ResNet-1D, which have been shown to perform well in biological sequence modeling tasks(Bileschi et al., 2022), offering a good trade-off between accuracy and computational efficiency compared to recurrent or attention-based models. Below is a brief introduction to each model:

i. AlexNet was a deep convolutional neural network (CNN) model specifically designed to handle one-dimensional data. The model combined multiple convolutional layers, pooling layers, and fully connected layers, along with ReLU activation functions and Dropout techniques, to prevent overfitting. AlexNet demonstrated outstanding performance in multi-label classification tasks, particularly in extracting temporal features.
ii. VGG was a classic deep CNN optimized for processing one-dimensional data. Unlike its traditional two-dimensional version, VGG used one-dimensional convolutional and pooling layers to learn temporal data features. By stacking multiple small convolutional kernels, VGG increased the depth of the network, enabling it to efficiently extract complex temporal features and perform excellently in multi-label classification tasks. The VGG16 model was employed in this study.
iii. GoogleNet was based on the Inception module, which used a combination of multiple convolutional kernels of different sizes within the same layer to extract multi-scale features. The model reduced computational complexity through global average pooling and efficiently handled one-dimensional data. In multi-label classification tasks, GoogleNet exhibited strong feature extraction capabilities and efficient computational performance.
iv. ResNet was a one-dimensional network structure based on residual networks (ResNet), which effectively addressed the gradient vanishing problem in deep networks by using one-dimensional convolutional layers and residual connections (skip connections). To prevent overfitting, ResNet added a Dropout layer to the original structure and used larger convolution kernels (e.g., 7×1 convolutions). This design enhanced the model’s generalization ability and accuracy, making it perform well in multi-label classification tasks. The ResNet18 model was employed in this study.
v. BERT was a pre-trained language model based on the Transformer architecture, specifically designed for processing one-dimensional sequence data. Using a bidirectional encoder, BERT effectively captured contextual information, allowing for better understanding of the sequential relationships in one-dimensional input data. BERT demonstrated strong text comprehension abilities in multi-label classification tasks, particularly suited for feature learning in high-dimensional data.
vi. ViT was a model that applied the Transformer architecture to one-dimensional data. It divided the input sequence into multiple small segments (patches) and utilized self-attention mechanisms to model long-range dependencies, allowing it to efficiently handle long sequence data. In multi-label classification tasks, ViT performed excellently, especially in handling complex temporal data with strong advantages.
vii. Mamba2 combined CNN with self-attention mechanisms to effectively handle complex label relationships and high-dimensional data. Through deep feature extraction and contextual modeling, Mamba2 demonstrated strong learning capabilities and high generalization ability, making it particularly suitable for multi-label classification tasks.

### 2.4 Experimental Validation

To evaluate the effectiveness of the expanded metal resistance gene library and the predictive performance of the classification model, we selected four representative genes for experimental validation. Two of the genes were derived from the expanded library but were not included in the BacMet2_EXP database (Table S3), while the other two were metal resistance genes identified from *Aeromonas* sp. NJAU223 (Chen et al., 2025) that were not part of the expanded library but were predicted by our model to confer metal resistance (Table S3). These four genes included the fpvA gene, which is associated with resistance to copper (Cu) and zinc (Zn), and the zntA gene, which is associated with resistance to cadmium (Cd) and lead (Pb). These genes were synthesized by Tsingke Biotechnology Co., Ltd., and were individually cloned into the pME6032 plasmid using seamless cloning. The recombinant plasmids were then transformed into *Escherichia coli* DH5α, generating four engineered strains: *E. coli* DH5α-O1, *E. coli* DH5α-O2, *E. coli* DH5α-O3, and *E. coli* DH5α-O4. The expression of each gene was confirmed using real-time quantitative PCR (Text S2). Based on the predicted metal resistance profiles, *E. coli* DH5α-O1 and DH5α-O3 were tested for their resistance to and removal of Cu and Zn, while *E. coli* DH5α-O2 and DH5α-O4 were assessed for Cd and Pb resistance and removal (Text S4). All experiments were performed in triplicate. Data were presented as means ± standard deviation (SD), and statistical significance was determined using one-way analysis of variance (ANOVA).

## 3. Results

### 3.1 Distribution, Sequence Characteristics, and Metal Associations of Resistance Genes in Environmental Microbes

Fig. 2a illustrated the distribution of the number of metal resistance genes across various metallic elements. Statistical analysis revealed that Zn and Cu exhibited the highest number of resistance genes, with both exceeding 250,000—significantly higher than the resistance gene counts of other metals such as nickel (Ni, approximately 175000 resistance genes), iron (Fe, approximately 165000 resistance genes), and cobalt (Co, approximately 135000 resistance genes). This finding suggested that resistance mechanisms related to Zn and Cu were highly representative in the genome, likely due to their widespread environmental distribution and biological significance. Additionally, the resistance gene numbers for Ni and Fe were also considerable, surpassing 200,000 and 150,000, respectively, reflecting the critical roles these metals play in microbial metabolism and enzyme activity. The resistance gene counts for Co, tungsten (W), and Cd were moderate, indicating their importance in specific ecological niches. In contrast, the resistance gene numbers for aluminum (Al), gold (Au), and bismuth (Bi) were relatively low, each below 10,000. For aluminum, despite its abundance in nature, the limited number of resistance genes may reflect both its lower bioavailability and potentially less research attention compared to other metals. In the case of gold and bismuth, their lower resistance gene numbers could be attributed to their relatively scarce natural occurrence, limited direct interactions with organisms, as well as possibly fewer studies focusing on their resistance mechanisms.

**Fig. 2.**
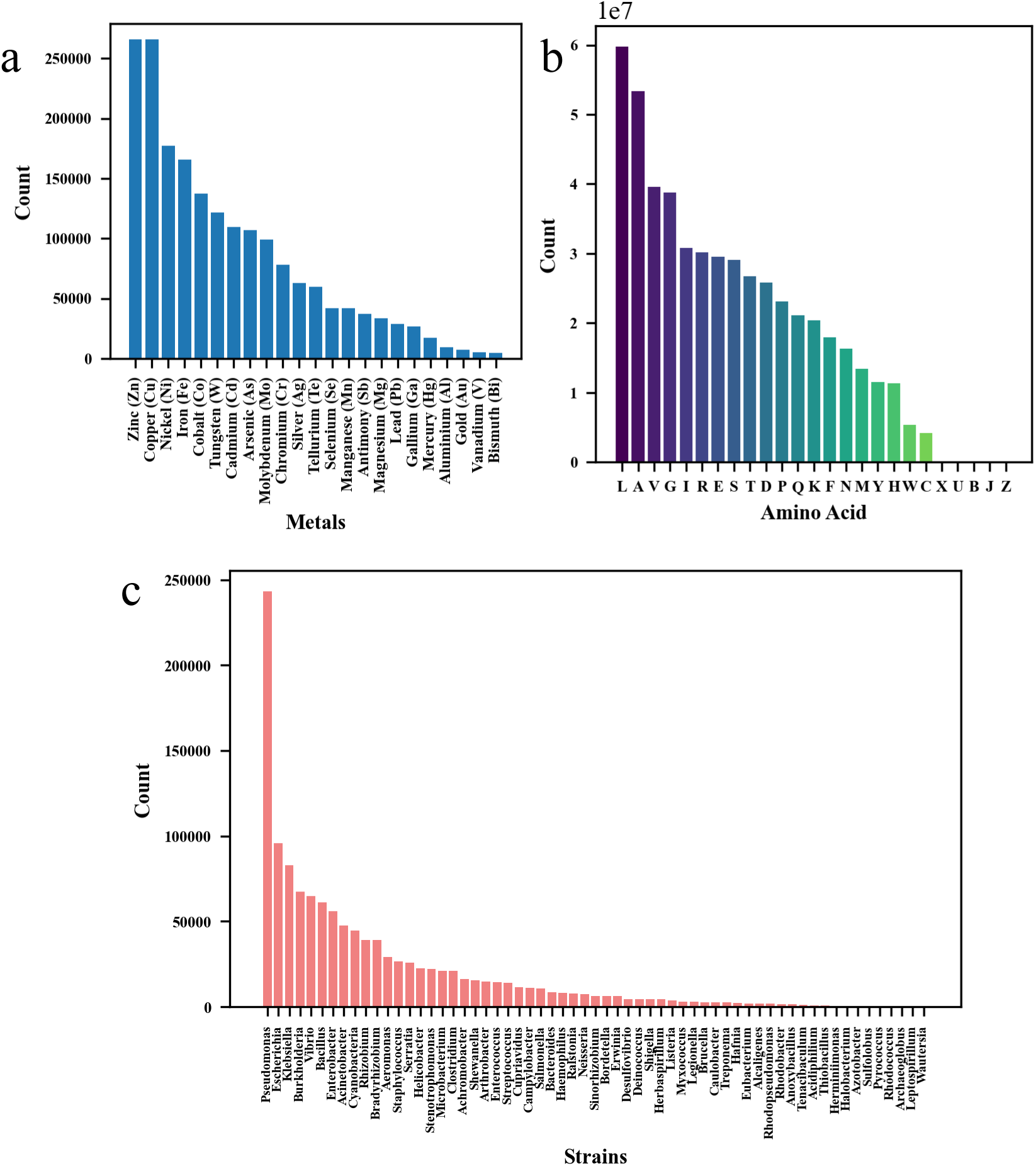
(a) Statistics of the number of metal resistance amino acid sequences. (b) Statistics of the amino acid counts in metal resistance sequences. (c) Statistics of the number of 64 prokaryotic species.

There were significant differences in the protein sequence numbers across various strains, with *Pseudomonas* possessing the highest number of sequences, approaching 250,000(Fig. 2c). This reflected the robust genetic reservoir of *Pseudomonas* in metal-contaminated environments, likely linked to its extensive ecological adaptability and its significant role in industrial, clinical, and environmental settings, which explained its focus in research. Following this, *Bacillus* and *Enterobacter* had approximately 100,000 and 80,000 protein sequences, respectively, indicating their high representation in environmental microbial communities. Other strains, such as *Bacillus thuringiensis* and *Acinetobacter*, had fewer than 10,000 sequences, likely due to lower research attention or less genomic data available in databases.

In terms of amino acid composition, Leucine (L) was the most abundant, exceeding 6×10⁷, significantly higher than other amino acids, suggesting its potential crucial structural or functional role in metal resistance-related proteins (Fig 2b). The next most abundant amino acids were alanine (A) and valine (V), with quantities approximately 5×10⁷ and 4.5×10⁷, respectively. As hydrophobic amino acids, these residues were likely associated with the hydrophobic core and stability of proteins. Glycine (G), Isoleucine (I), Arginine (R), and Glutamic acid (E) exhibited quantities ranging from 2×10⁷ to 3×10⁷, possibly playing important roles in metal binding, enzyme activity, or protein folding. Tryptophan (W), Cysteine (C), Selenocysteine (U), and other rare amino acids (e.g., unknown or unspecified amino acid (X), Asparagine or Aspartic (B), Leucine or Isoleucine (J), Glutamine or Glutamic (Z)) had low frequencies of occurrence, with quantities all below 5×10⁶.

The amino acid sequence lengths were primarily concentrated between 200 and 600(Fig 3a), with a right-skewed distribution, indicating that most sequences were relatively short, though a substantial number of longer sequences (>1000) were present. The normal distribution fit showed that the average sequence length was 416.88, with a standard deviation of 235.82. The violin plots for different strains further confirmed this trend: the median sequence length for most strains was around 400, with a distribution concentrated between 200 and 600(Fig. S1a), suggesting that the high-frequency region of the overall distribution was widely representative among the strains. However, some strains, such as *Pseudomonas* and *Bacillus*, exhibited greater sequence length variability (>1000), whereas others, such as *Achromobacter* and *Rhodococcus*, showed more concentrated distributions, highlighting the diversity in sequence lengths of metal resistance proteins across strains.

**Fig. 3.**
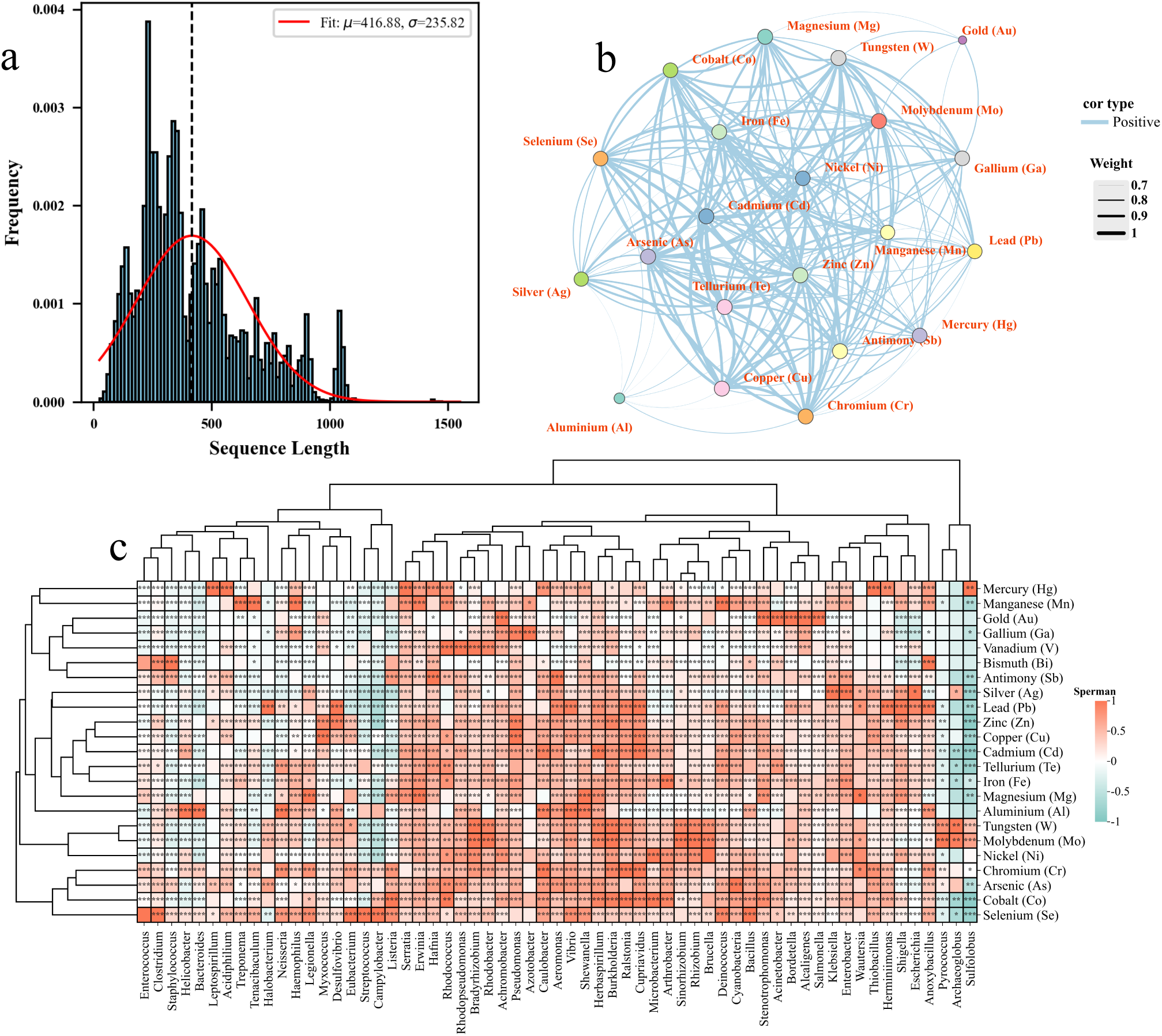
(a) Statistics of the lengths of metal resistance amino acid sequences. (b) Metal-metal network based on metal resistance amino acid sequences. (c) Metal-species correlation analysis based on metal resistance amino acid sequences. *: p < 0.05; **: p < 0.01; ***: p < 0.001.

Overall, the sequence lengths of different metal resistance-related proteins were predominantly concentrated between 200 and 600(Fig. S1b), consistent with the overall length frequency distribution. The median length of most metal resistance sequences was close to 400, indicating that the overall distribution trend was similar across metal categories. However, metals such as Pb, chromium (Cr), and Cd exhibited a higher frequency of longer sequences, whereas less common metals, including Au, Bi, and Ga, showed a narrower and shorter amino acid length distribution. Similar to *Pseudomonas* and other strains with significant length variability (Fig. S1a), the violin plots for metals such as Zn, Cu, and Fe also showed broader distribution ranges, suggesting that these metal resistance proteins could be contributed by various strains and involve diverse sequence length requirements.

The Fig.S2c indicated that *Pseudomonas* and *Escherichia* were dominant strains associated with resistance to multiple metals, with their numbers significantly higher than those of other strains, indicating their extensive genetic reservoirs and adaptability in metal resistance. Significance analysis further demonstrated that these strains exhibited strong positive correlations with metals such as Zn, Cu, Ni, Fe, and Pb, reflecting their crucial adaptability in multi-metal contaminated environments (Fig. 3c). Furthermore, specific strains such as *Rhodobacter* and *Bradyrhizobium* showed strong correlations with Ni and molybdenum (Mo), suggesting that these strains may possess specificity for certain metal resistance functions. Regarding amino acids, Leucine, Alanine, Valine, and Glycine were the primary amino acids in metal resistance proteins (Fig. S2b), showing significant positive correlations with several metals (e.g., Zn, Cu, Ni), consistent with their importance in protein structural stability and functional execution (Fig. S2c). Specific amino acids, such as Cysteine, exhibited strong correlations with mercury (Hg) and Cd, potentially playing a role in binding and detoxification through thiol groups, while tryptophan and methionine showed higher correlations with Zn and Cu, suggesting their key roles in specific metal resistance functions. Moreover, Zn, Cu, Ni, and Fe formed a strong correlation cluster (Fig 3b), suggesting that resistance to these metals may involve synergistic mechanisms, whereas Cd and Pb, which often co-exist in highly polluted environments, displayed a high correlation in their resistance. In contrast, rare metals (e.g., Au, Bi, Ga) had fewer resistance proteins and lower correlations (Fig 3b), likely due to their mechanisms not being fully explored.

### 3.2 Optimization of Deep Learning Models for Metal Resistance Gene Classification

To evaluate the robustness and generalization ability of each model in the multi-label classification task of metal resistance genes, we compared the overall Jaccard score of each model on both the training and validation datasets. The best-performing models were identified based on their overfitting risk, defined as the relative difference in overall Jaccard score between the training and validation sets.

Both ResNet and VGG demonstrated strong performance and robustness (Fig. 4a). However, a further comparison of their performance on the training and validation datasets revealed that ResNet’s overall performance slightly outperformed that of VGG. Specifically, ResNet achieved Jaccard indices of 98.92% and 98.94% on the training and validation datasets (Fig. 4b), respectively, with a near-perfect match (overfitting risk of −0.02%). This reflects ResNet’s more thorough learning during training and its exceptional generalization ability, achieving near-identical performance on the validation set compared to the training set. In contrast, VGG exhibited a slightly higher Jaccard score on the validation set (99.18%); however, the Jaccard score on the training set (99.74%) showed an overfitting risk of 0.56%, indicating mild overfitting during training. Therefore, ResNet demonstrated higher robustness and generalization ability, with consistent performance between the training and validation datasets and a near-zero overfitting risk, making it the superior model compared to VGG.

**Fig. 4.**
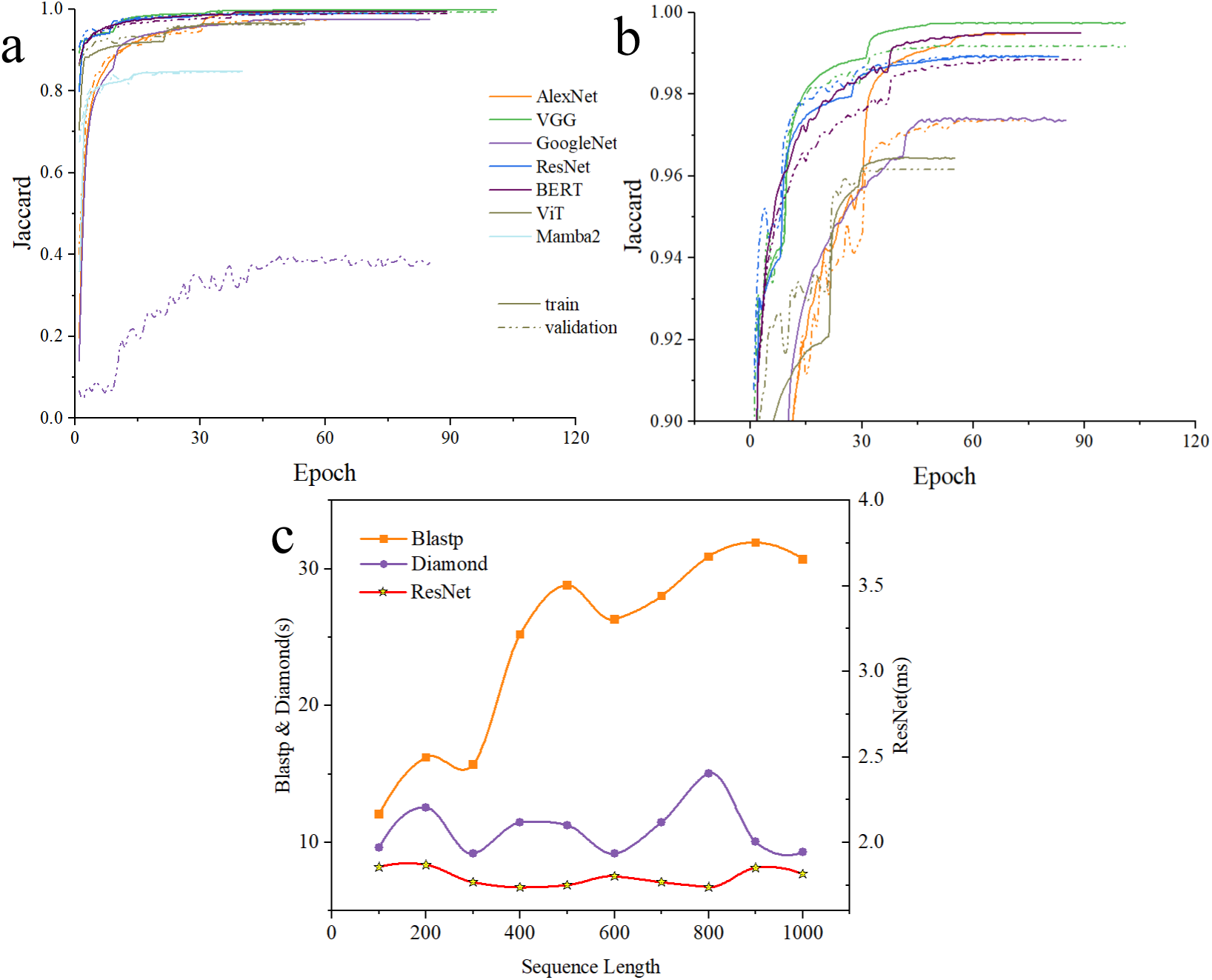
(a) Jaccard scores of the AlexNet, VGG, GoogleNet, ResNet, BERT, ViT, and Mamba2 models as a function of Epoch. (b) A magnified view of Fig. (a). (c) Comparison of runtime for different lengths of amino acid sequences on Blastp, Diamond software, and the ResNet model. CPU: i7-11700, GPU: RTX-3060.

BERT achieved a Jaccard score of 98.83% on the validation set and 99.49% on the training set, with an overfitting risk of 0.66% (Fig. 4b). While its performance on the validation set was slightly lower than that of ResNet and VGG, it still performed excellently and demonstrated strong robustness in multi-label classification tasks. In contrast, the performance of AlexNet and GoogleNet was relatively weaker. AlexNet achieved a Jaccard score of 97.35% on the validation set and 99.49% on the training set, with an overfitting risk of 2.15%, suggesting limited generalization ability and potential difficulty in handling complex tasks. GoogleNet’s Jaccard score on the validation set was only 39.79%, significantly lower than the training set’s 97.43%, with an overfitting risk of 59.16%, indicating poor adaptability for multi-label classification tasks. As for ViT and Mamba2, both performed below 97% on the training and validation datasets, suggesting that further optimization of the model architecture was required. Based on the above analysis, ResNet was selected as one of the top-performing models for this task, demonstrating exceptional robustness and generalization ability with a high Jaccard score, near-zero overfitting risk, and consistency between training and validation performance. VGG ranked second, with slightly higher validation set performance but exhibiting mild overfitting. BERT, as the third-best model, is suitable for specific scenarios. In contrast, the performance of AlexNet, GoogleNet, ViT, and Mamba2 was insufficient, with poor generalization ability, making them unsuitable as the primary model choices.

We performed a detailed analysis of the F1 Macro, Precision, Recall, and Hamming Loss metrics for each model in the multi-label classification task. On the training set, ResNet achieved an F1 Macro of 99.44% (Fig. S3b), Precision of 99.81% (Fig. S3c), Recall of 98.97% (Fig. S3d), and Hamming Loss of 0.09% (Fig. S3a), ranking just below VGG and BERT. Despite some slightly lower metrics, ResNet’s consistency between the training and validation datasets highlighted its stability during training. On the validation set, ResNet achieved an F1 Macro of 99.33%, Precision of 99.66%, Recall of 99.01%, and Hamming Loss of 0.10%, outperforming BERT and AlexNet, and approaching VGG, demonstrating exceptional generalization ability and robustness (Fig. S3). In contrast, GoogleNet performed poorly on the validation set, with an F1 Macro of only 59.13% and a Hamming Loss of 5.32%.

Prior to model training, we applied preprocessing methods such as removing low-frequency amino acids, random substitution, oversampling, and undersampling to enhance data quality and balance the sample distribution. The test results for the ResNet model showed that the highest Jaccard score, 98.91%, was achieved when all preprocessing methods were applied (Table 1). After removing low-frequency amino acid removal or random substitution, the Jaccard score dropped to 98.77% and 96.56%, respectively. Further removal of oversampling led to a decrease to 93.52%, indicating that each preprocessing step was crucial for improving model performance (Table 1).

**Table 1.** Effect of preprocessing on the Jaccard scores of the test set for the ResNet model.

| Preprocessing | Test-Jaccard (%) |
| --- | --- |
| **ResNet** | **98.91** |
| -unwant | 98.77 |
| -substitution | 96.56 |
| -substitution-oversample | 93.52 |

**Note:** -unwant: No low-frequency amino acids were removed. -substitution: No random substitution was performed. -substitution-oversample: No random substitution and oversampling was performed.

We evaluated the impact of amino acid augmentation methods, including deletion, swap, insertion, and substitution, on the ResNet model. When substitution was applied alone, the Jaccard score reached a peak of 98.91%, and combinations with swapping and insertion methods (such as swa+sub, ins+sub) also performed excellently (Table 2). Combinations that included the deletion method significantly reduced performance, and when all augmentation methods were applied together, the Jaccard score dropped to 96.71%, suggesting that an excessive number of augmentation strategies might introduce noise.

**Table 2.**
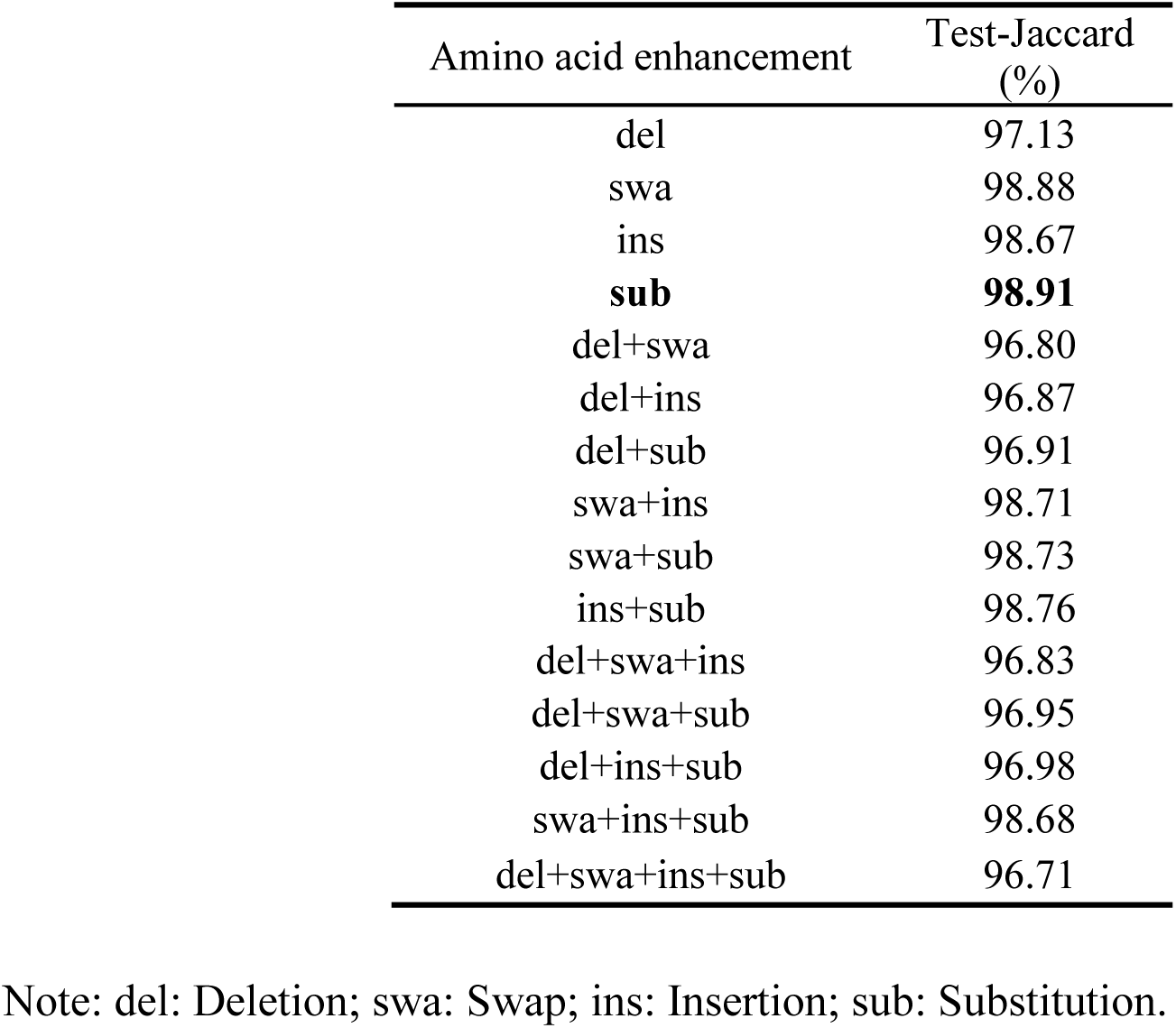
Effect of random deletion, swap, insertion, and substitution on the Jaccard scores of the test set for the ResNet model.

| Amino acid enhancement | Test-Jaccard (%) |
| --- | --- |
| del | 97.13 |
| swa | 98.88 |
| ins | 98.67 |
| **sub** | **98.91** |
| del+swa | 96.80 |
| del+ins | 96.87 |
| del+sub | 96.91 |
| swa+ins | 98.71 |
| swa+sub | 98.73 |
| ins+sub | 98.76 |
| del+swa+ins | 96.83 |
| del+swa+sub | 96.95 |
| del+ins+sub | 96.98 |
| swa+ins+sub | 98.68 |
| del+swa+ins+sub | 96.71 |

Note: del: Deletion; swa: Swap; ins: Insertion; sub: Substitution.

In the comparison of embedding methods, nn.embedding achieved the highest Jaccard score on the test set, 98.91%, significantly outperforming One-Hot (98.80%) and Protvec (98.77%), while ESM-1b and TAPE performed relatively weaker (96.47% and 95.78%, respectively) (Table 3). This suggests that feature learning based on embedding layers is more effective in capturing the contextual information and underlying semantic features of amino acid sequences.

**Table 3.**
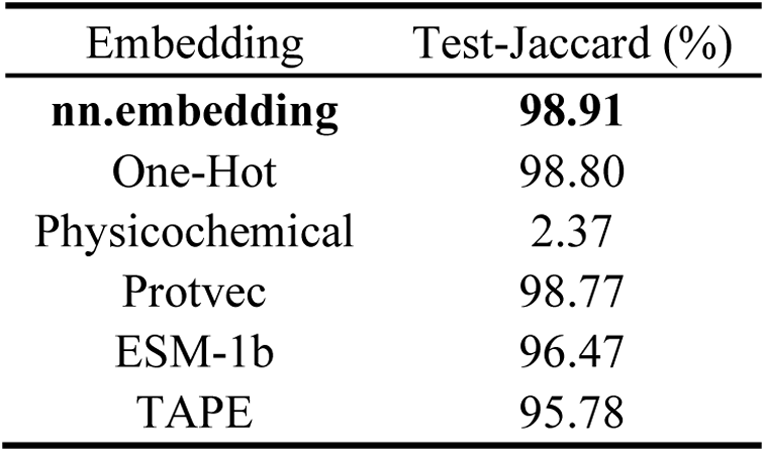
Effect of different embedding methods on the Jaccard scores of the test set for the ResNet model.

| Embedding | Test-Jaccard (%) |
| --- | --- |
| **nn.embedding** | **98.91** |
| One-Hot | 98.80 |
| Physicochemical | 2.37 |
| Protvec | 98.77 |
| ESM-1b | 96.47 |
| TAPE | 95.78 |

A comprehensive evaluation of the model parameters, file size, Jaccard score, and training time revealed that while VGG achieved the highest Jaccard score (99.15%), it had the largest number of parameters, file size, and longest training time. ResNet, with fewer parameters (8.79M), smaller file size (34.28Mb), and faster training speed (475.83s/epoch), achieved a good balance between performance and resource efficiency (Table 4). BERT, although resource-efficient (3.75M parameters, 12.88Mb file size), had longer training times (6041.11s/epoch). ViT and GoogleNet performed poorly in terms of training time and performance, while Mamba2, despite having very low resource consumption, achieved a Jaccard score of only 84.82%, making it unsuitable for complex tasks.

**Table 4.**
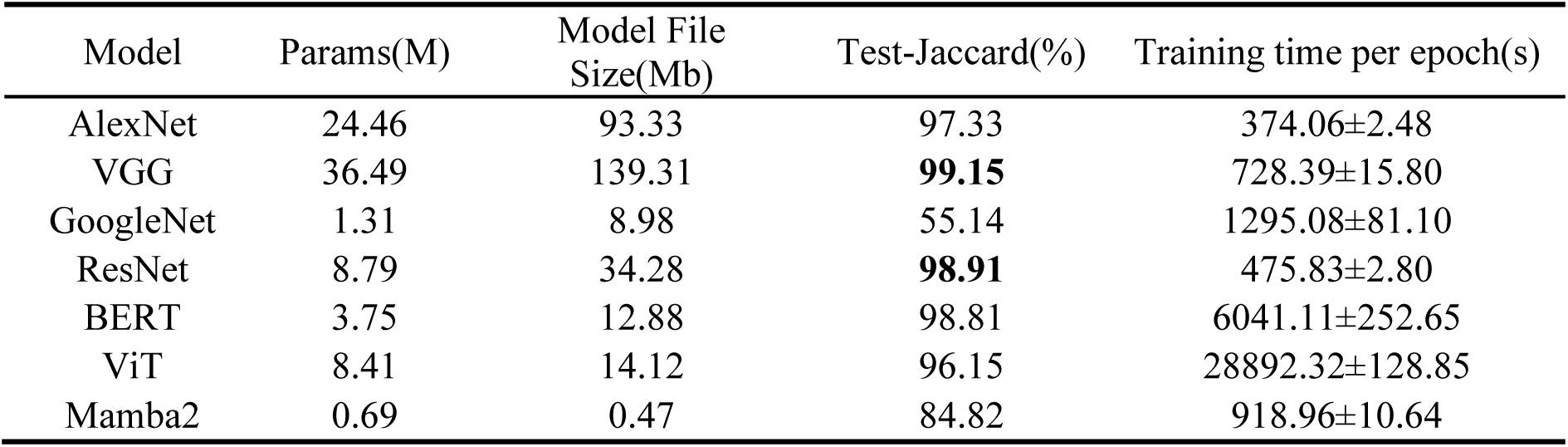
Comparison of parameters, file size, Jaccard scores on the test set, and training time for different models.

| Model | Params(M) | Model File Size(Mb) | Test-Jaccard(%) | Training time per epoch(s) |
| --- | --- | --- | --- | --- |
| AlexNet | 24.46 | 93.33 | 97.33 | 374.06±2.48 |
| VGG | 36.49 | 139.31 | **99.15** | 728.39±15.80 |
| GoogleNet | 1.31 | 8.98 | 55.14 | 1295.08±81.10 |
| ResNet | 8.79 | 34.28 | **98.91** | 475.83±2.80 |
| BERT | 3.75 | 12.88 | 98.81 | 6041.11±252.65 |
| ViT | 8.41 | 14.12 | 96.15 | 28892.32±128.85 |
| Mamba2 | 0.69 | 0.47 | 84.82 | 918.96±10.64 |

Through t-SNE dimensionality reduction analysis, the untreated amino acid sequences did not exhibit clear clustering, while after processing with ResNet, distinct clusters of different metal resistance categories were formed, demonstrating that the model effectively extracted discriminative features (Fig. 5). The model achieved precision, recall, and F1 scores greater than 99% for metals such as Al, selenium (Se), and magnesium (Mg), demonstrating excellent performance (Table S1). However, the F1 scores for metals such as Cu, Au, silver (Ag), and Zn were lower (0.62-0.88), indicating the need for further optimization. Additionally, the F1 scores for Cd and Hg were 0.91 and 0.90, respectively, showing that the model still has room for improvement in handling complex metal resistance mechanisms (Table S1).

**Fig. 5.**
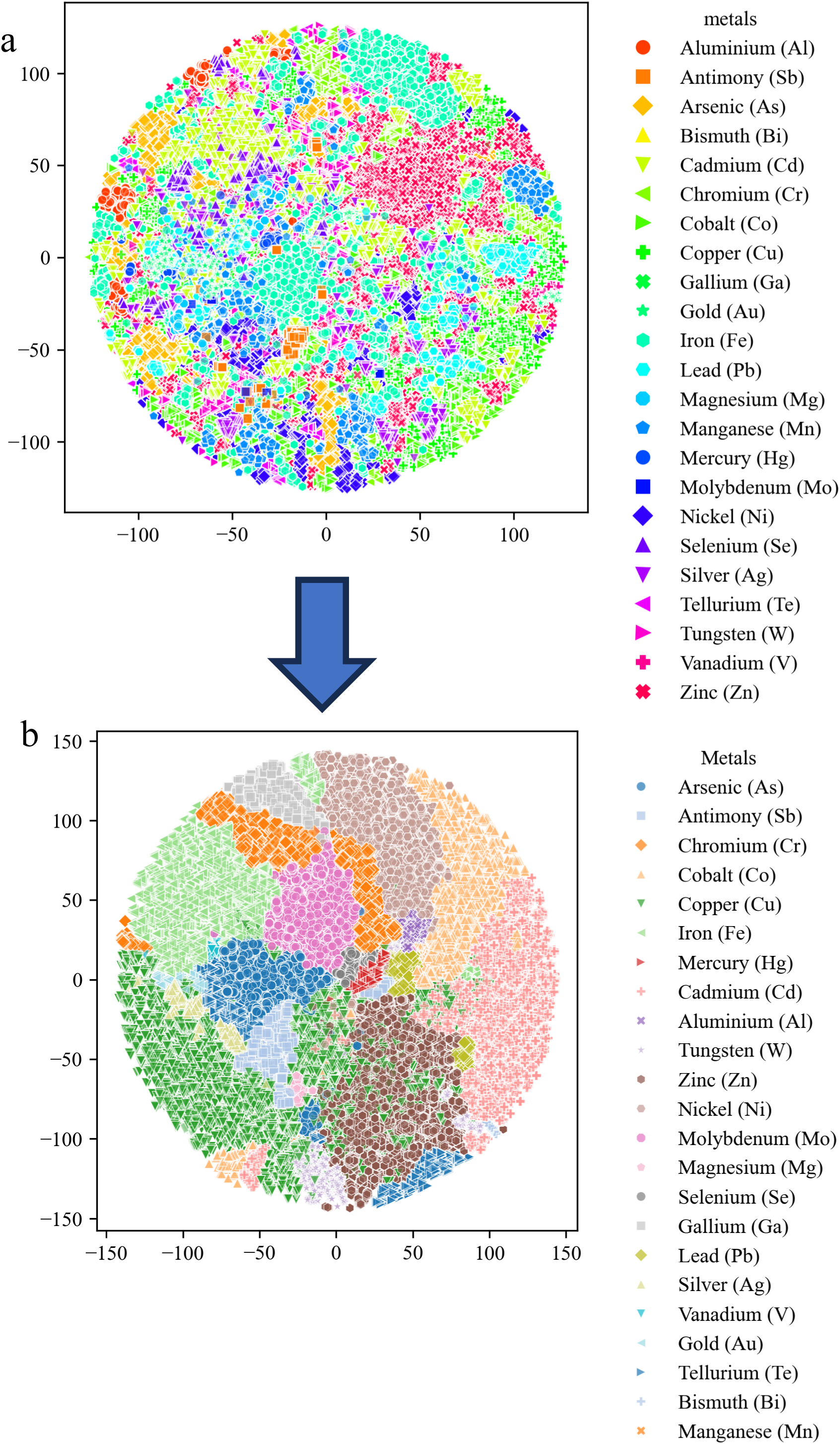
(a) t-SNE dimensionality reduction of metal resistance amino acid sequences. (b) t-SNE dimensionality reduction of metal resistance amino acid sequences processed through the ResNet model.

To further validate the practicality and efficiency of the top-performing model, ResNet, we compared its prediction time with that of traditional sequence alignment methods, BLASTp and DIAMOND. The experiment measured the prediction times of the three methods on sequences of varying lengths (100-1000 bp). The results demonstrated that ResNet had a significant advantage in prediction speed. For shorter sequences (100 bp), the prediction times of BLASTp and DIAMOND were 12.07s and 9.63s, respectively, while ResNet’s prediction time was only 1.86ms, approximately 5000 times faster than DIAMOND and 6500 times faster than BLASTp (Fig. 4c). As the sequence length increased, the prediction times for BLASTp and DIAMOND gradually grew, reaching 30.68s and 9.27s, respectively, for 1000 bp sequences, while ResNet’s prediction time remained at 1.81ms, maintaining a high speed advantage. The experimental results showed that ResNet’s prediction time was nearly constant across varying sequence lengths, remaining stable in the range of 1.73-1.86ms, demonstrating exceptional robustness to sequence length. This characteristic significantly outperformed BLASTp and DIAMOND, with BLASTp’s prediction time increasing non-linearly with sequence length. Additionally, the efficiency of ResNet is attributed to the deep learning model’s ability to map input sequences to label categories efficiently using pre-trained parameters, rather than performing one-by-one sequence alignments. As shown in Table 5, the ResNet model significantly outperformed both BLASTp and DIAMOND across all evaluation metrics. In particular, ResNet achieved a Jaccard score of 98.91%, compared to 98.19% for BLASTp and 98.16% for DIAMOND, demonstrating its superior predictive performance. In conclusion, ResNet, with its extremely low prediction time and stability across sequence lengths, significantly outperforms traditional sequence alignment methods such as BLASTp and DIAMOND, making it especially suitable for high-throughput biological sequence analysis tasks that require fast processing.

**Table 5.** Comparison of predictive performance between ResNet, BLASTp, and DIAMOND.

| Method | Jaccard(%) | F1 Macro(%) | Precision(%) | Recall(%) |
| --- | --- | --- | --- | --- |
| ResNet | 98.91 | 99.33 | 99.65 | 99.00 |
| BLASTp | 98.19 | 98.61 | 98.63 | 98.58 |
| DIAMOND | 98.16 | 98.60 | 98.59 | 98.60 |

### 3.3 Experimental Validation

To further validate the expanded metal resistance gene library and the predictive performance of the model, metal resistance and removal experiments were conducted by overexpressing the selected genes. As shown in Fig. S4a, agarose gel electrophoresis confirmed the successful insertion of all four gene fragments. Fig. S4b demonstrated that the expression levels of these genes were significantly higher than those of the control group (*p* < 0.05).

In terms of metal resistance, *E. coli* DH5α-O1 and DH5α-O3 exhibited no significant growth difference compared to *E. coli* DH5α in the absence of Cu or Zn. However, under treatment with 5 mg/L Cu or Zn, the growth of *E. coli* DH5α was significantly lower than that of *E. coli* DH5α-O1 and DH5α-O3 (Fig S5). Similarly, *E. coli* DH5α-O2 and DH5α-O4 showed comparable growth to *E. coli* DH5α without Cd or Pb treatment, while exposure to 2 and 5 mg/L Cd or 5 mg/L Pb significantly inhibited the growth of *E. coli* DH5α compared to that of *E. coli* DH5α-O2 and DH5α-O4 (Fig S5).

Regarding heavy metal removal efficiency, *E. coli* DH5α-O1 and DH5α-O3 showed significantly higher Cu removal at 0.5–2 mg/L and Zn removal at 0.5 and 1 mg/L compared to *E. coli* DH5α (Fig S6). However, this difference diminished as the concentrations of Cu and Zn increased. Similarly, *E. coli* DH5α-O2 and DH5α-O4 exhibited significantly higher Cd removal at 0.5–2 mg/L and Pb removal at 0.5 and 1 mg/L than *E. coli* DH5α, but the enhancement weakened with increasing Cd and Pb concentrations (Fig S6).

In summary, overexpression of metal resistance genes enhanced both the heavy metal tolerance and removal capabilities of *E. coli*, thereby successfully validating the reliability of the expanded metal resistance gene library and the predictive power of the classification model.

## 4. Discussion

Heavy metal pollution continues to be a critical global challenge, threatening both ecosystems and human health(Das et al., 2023). Understanding the genetic basis of microbial resistance is paramount for devising effective remediation strategies, as metal resistance genes enable microorganisms to survive in contaminated environments and mitigate toxicity(Das et al., 2016). Building on earlier work with the BacMet database(Pal et al., 2014), we substantially expanded its coverage from only 753 experimentally verified sequences to a total of 1,219,137 unique amino acid sequences. This enrichment addresses the longstanding limitation of slow database updates and restricted data scope, thus providing an up-to-date, large-scale resource for the identification and classification of metal resistance genes. Additionally, we implemented seven deep learning models—including AlexNet, VGG, GoogleNet, ResNet, BERT, ViT, and Mamba2—to conduct multi-label classification. Our results reveal that ResNet achieves remarkably high predictive accuracy, with a Jaccard score of 98.91%. Additionally, it provides annotation speeds that are substantially faster than traditional sequence alignment methods, such as BLASTp and DIAMOND. These findings underscore both the importance of expanding metal resistance gene repositories and the potential of advanced deep learning algorithms to accelerate and refine genomic annotations, thereby laying a robust foundation for future bioremediation efforts (Bileschi et al., 2022; Brandes et al., 2022; He et al., 2016).

Compared to other studies, this research not only expanded the metal resistance amino acid sequence database but also revealed the relationships between different species, metal resistance amino acid sequences, and their amino acid compositions through statistical analyses, offering new insights into microbial adaptation mechanisms in metal-polluted environments. This aligns with Wu et al.’s study, which predicted quorum sensing (QS) molecules in the human gut microbiome using machine learning methods, but our study advanced further in terms of data scale and model complexity, providing a more detailed and efficient solution(Wu et al., 2022).

These findings were consistent with previous studies on the genus *Pseudomonas*, which demonstrated strong genetic reservoir capabilities in metal-polluted environments (Fig. S2c), further supporting our results(Fakhar et al., 2022). *Pseudomonas* species were ubiquitous in various environmental media, mainly due to their broad metabolic range(Hesse et al., 2018), which enabled them to degrade a variety of organic and inorganic compounds(Hamamura et al., 2013), and their strong resistance to antibiotics, heavy metals, detergents, and organic solvents(Ansari and Malik, 2007; Haritash and Kaushik, 2009; Pardo et al., 2003). *Pseudomonas* reduced heavy metal toxicity through various mechanisms, such as transformation, volatilization, fixation, complexation, and oxidation(Dixit et al., 2015). Notably, *Pseudomonas* showed significant tolerance to Cu and Zn, with *Pseudomonas koreensis* demonstrating resistance to Zn up to 3000 ppm(Khatoon et al., 2020), which was attributed to its abundance of Cu and Zn resistance genes (Fig. 3c). The absorption of Ni and Mo by *Bradyrhizobium* was primarily related to its symbiotic activity with legumes(Johnston et al., 2001). On the other hand, *Rhodobacter* required Fe, Mg, Ca, and Ni ions for hydrogen production and nitrogen fixation(Gabrielyan et al., 2016). The high frequency of Leucine, Alanine, Valine, and Glycine in metal resistance proteins in the study was related to their ability to form stable complexes with metal ions such as Cu and Zn(Fig. 1c)(Hamadi et al., 2018). Cysteine could form similar complexes with Cd and Hg, such as Hg-cysteine and Cd-cysteine complexes(Gruff and Koch, 1990). Microorganisms toxic to Cu and Zn may secrete more tryptophan to detoxify these metals (Li et al., 2023; Ye et al., 2023).

A key innovation in ResNet for model prediction was the introduction of skip connections, where signals were not only transmitted sequentially between network layers but could also skip certain layers for direct connections. This design aimed to address the degradation problem during deep neural network training, where accuracy decreased as the number of layers increased (Glorot et al., 2011; Zilly et al., 2017). Although AlexNet and VGG networks were relatively shallow, the lack of residual connections meant that adding more layers did not improve performance and might even lead to overfitting (Fig. 4b). In many standard image classification tasks (such as ImageNet)(Krizhevsky et al., 2017), ResNet-18 typically outperformed AlexNet, VGG, GoogleNet, and others in terms of accuracy(He et al., 2016). Amino acid sequences were essentially structured sequential data, where each amino acid has a specific relationship with neighboring amino acids. Therefore, CNNs, such as ResNet-18, could effectively extract local features (such as amino acid proximity, local structures, etc.) through convolution operations. In such tasks, ResNet captured effective features through local convolutional layers, leading to better classification performance. BERT and ViT models, based on self-attention mechanisms, excel at capturing long-range dependencies, but they were not as effective as CNNs in structured sequential data, especially in tasks where short sequences or local features were critical. While VGG and BERT achieved higher Jaccard scores, their slightly elevated overfitting risks suggest reduced robustness. ResNet offered a better trade-off between accuracy and generalization, making it more suitable for stable deployment in metal resistance prediction. In the metal resistance amino acid sequence database we constructed, the longest amino acid sequence was 1500 bp, while the majority of sequences were shorter than 1000 bp, making CNNs more suitable for classifying metal resistance amino acid sequences compared to self-attention mechanisms. Bileschi et al. also employed convolutional neural networks with residual connections, achieving an accuracy of 98% in the Pfam multi-classification task(Bileschi et al., 2022).

In addition to the model itself, data preprocessing played a crucial role in enhancing the performance of the model(Domingos, 2012). The preprocessing methods used in this study have also proven effective in other fields. By removing low-frequency features to reduce noise contamination, Liu et al. enhanced model performance by eliminating low-frequency words, improving accuracy from 98.2% to 99.2%, demonstrating the effectiveness of noise removal in improving handwritten digit recognition accuracy(Liu and Zhang, 2012). For amino acid augmentation, methods such as rotation, translation, scaling, cropping, and flipping, which were commonly used in image augmentation, were applied, significantly improving model accuracy by 3% to 7%(Shorten and Khoshgoftaar, 2019). In time-series data, common augmentation techniques included temporal shifting, interpolation, noise addition, and random perturbation. Studies have shown that applying these methods increased the accuracy of time-series classification models by 5% to 10%(Wei and Zou, 2019). There were three primary reasons why these augmentation methods enhanced model performance: first, they increased data diversity. By simulating different input variations, the model became more robust when encountering unseen data. Second, they reduced overfitting. By augmenting the training dataset, especially in cases of small datasets or data imbalance, overfitting to the training data was minimized. Finally, they improved generalization. Augmentation methods helped the model better understand the diversity of the data, leading to more stable performance in real-world applications. Despite the improvements brought by preprocessing and zero-padding, we acknowledge their potential limitations. The current model may be sensitive to curated inputs, and fixed-length padding may not fully reflect biological structure. In future work, we will explore dynamic padding, attention-based models, and external dataset validation to enhance robustness and biological relevance.

GPU-accelerated methods based on deep learning, compared to traditional CPU-based computing, significantly increased computational speed, reduced training and inference time(Silvano et al., 2023), and improved model accuracy through automatic feature learning. Deep learning models could handle larger datasets, capture more complex patterns, and significantly improve efficiency through parallel computation(Krizhevsky et al., 2017; Simonyan, 2014). The BERT model(Devlin et al., 2019), in various NLP tasks (e.g., text classification, question answering), was faster and more accurate than traditional feature-based models (e.g., SVM, TF-IDF). Studies have shown that deep learning methods (e.g., deep neural networks) could achieve faster results than traditional methods (e.g., BLASTp) in DNA sequence alignment tasks, while maintaining high accuracy(Alipanahi et al., 2015). Although ResNet achieves faster inference compared to BLASTp and DIAMOND, its training process requires substantial time and computational resources. In contrast, alignment-based tools, while slower at runtime, do not require pre-training and can flexibly adjust speed-accuracy trade-offs via parameter tuning. These methods thus offer complementary strengths for different application scenarios. Our comparison further highlights the advantage of ResNet over classical alignment-based tools. While the Jaccard score achieved by BLASTp (98.22%)(Bileschi et al., 2022)aligns with our baseline results, our ResNet model slightly outperformed the previously reported ProtCNN model (98.72%), indicating both robustness and potential for further improvement through architecture optimization.

However, the model performed relatively poorly in classifying certain metals (e.g., Cu, Au, Ag), with F1 scores of 0.65, 0.62, and 0.71, indicating that the model’s recognition ability in these categories could be further improved. The model showed reduced performance for metals such as Cu, Au, and Ag, likely due to label imbalance, sequence variability, and cross-metal functional similarity. Secondly, although ResNet performed excellently overall, its recognition of resistance proteins for very rare metals (e.g., Au, Bi, Ga) remained limited, suggesting the need for further optimization of the model architecture or the addition of relevant data to improve classification performance. Future improvements may include metal-specific augmentation, adaptive loss functions, and biologically informed modeling strategies.

Unlike previous studies that focused solely on gene prediction using machine learning models(Wu et al., 2022), the present study further validated both the expanded metal resistance gene library and the predictive performance of the classification model. The genes inserted into *E. coli* DH5α-O1 and DH5α-O3 encoded outer membrane proteins capable of binding various metals(Hannauer et al., 2012), with sequence similarities of only 57.6% and 24.9% (Table S3), respectively, to entries in the BacMet2_EXP database. The genes introduced into *E. coli* DH5α-O2 and DH5α-O4 encoded P-type ATPases(Palmgren and Nissen, 2011), a class of membrane-associated transport proteins that utilize ATP hydrolysis to drive the transmembrane transport of ions or lipids, thereby facilitating the efflux of intracellular heavy metals. These transporters played a critical role in maintaining cellular homeostasis and were also found to be essential for *Aeromonas* sp. NJAU223 in resisting cadmium stress(Chen et al., 2025).

In summary, ResNet demonstrated significant advantages in multi-label classification tasks due to its stability, excellent generalization ability, and resource efficiency. Users could test the model via the PMRAAS website (https://s3.v100.vip:16165). To further validate the practical applicability and generalizability of the PMRAAS platform, we conducted a comparative analysis using protein sequences reported in previous studies (see Table S2). These sequences originated from species not included in our training dataset. The predictions generated by PMRAAS showed a high level of agreement with the experimentally validated metal resistance annotations, supporting the model’s ability to generalize beyond the training domain. This result highlights the robustness of our multi-label classification framework and the advantage conferred by the significantly expanded dataset of over 1.2 million sequences. Proper preprocessing and embedding methods further enhanced the model’s performance, but optimization was still needed for certain metal categories. Overall, the ResNet model significantly improved the separability of metal resistance features in amino acid sequences, and the clustering results after dimensionality reduction validated the model’s learning capability. Although the model showed excellent performance for most metals, further optimization is required for challenging metals (e.g., Cu, Au, and Ag) to improve classification accuracy and overall robustness. Although the model achieved strong overall performance, its accuracy on rare metals such as Au and Ag was limited due to data imbalance and sequence diversity. Moreover, generalization to unseen resistance genes outside the current dataset remains an open challenge. Future work will explore external validation, transfer learning, and cross-database modeling to improve the robustness and applicability of the model. These results provided important reference directions for future research and highlight the necessity of in-depth analysis of specific metal resistance mechanisms.

## 5. Conclusions

In this study, we significantly enhanced the BacMet database by expanding it into a comprehensive resource for metal resistance genes. Leveraging the superior performance of the ResNet model, our approach achieved exceptional accuracy and robustness in multi-label classification tasks, outperforming traditional sequence alignment methods. The predictive capability of the expanded gene library and the model was further validated through physiological experiments Moreover, the computational efficiency of ResNet enables unparalleled prediction speed, rendering it highly suitable for high-throughput analyses of biological sequences. To facilitate practical applications, we developed the PMRAAS website, which offers rapid and accurate online predictions of metal resistance, thereby supporting both research and remediation efforts. Collectively, these advancements provide valuable insights into the mechanisms underlying microbial adaptation to heavy metal contamination and lay a robust foundation for future bioremediation strategies. Nonetheless, limitations remain in predicting resistance to rare metals such as Au and Ag, due to data scarcity and functional overlap. Future work will focus on expanding the curated database, enhancing model robustness through external validation, and improving predictive performance on underrepresented categories.

## ACKNOWLEDGEMENT

The work was supported by the National Natural Science Foundation of China (32171623, 31770548).

